# Unsupervised machine learning reveals temporal components of gene expression in HeLa cells following release from cell cycle arrest

**DOI:** 10.1101/2022.07.13.499875

**Authors:** Tom Maimon, Yaron Trink, Jacob Goldberger, Tomer Kalisky

**Author notes:** Correspondence to: Tomer Kalisky, Faculty of engineering, Bar-Ilan University, Ramat Gan, Israel 52900.

## Abstract

Gene expression measurements of tissues, tumors, or cell lines taken over multiple time points are valuable for describing dynamic biological phenomena such as the response to growth factors. However, such phenomena typically involve multiple biological processes occurring in parallel, making it difficult to identify and discern their respective contributions at any time point. Here, we demonstrate the use of unsupervised machine learning to deconvolve a series of time-dependent gene expression measurements into its underlying temporal components. We first downloaded publicly available RNAseq data obtained from synchronized HeLa cells at consecutive time points following release from cell cycle arrest. Then, we used Fourier analysis and Topic modeling to reveal three underlying components and their relative contributions at each time point. We identified two temporal components with oscillatory behavior, corresponding to the G1-S and G2-M phases of the cell cycle, and a third component with a transient expression pattern, associated with the immediate-early response gene program, regulation of cell proliferation, and cervical cancer. This study demonstrates the use of unsupervised machine learning to identify hidden temporal components in biological systems, with potential applications to early detection and monitoring of diseases and recovery processes.

## INTRODUCTION

Complex biological systems such as tissues and tumors contain millions of cells. These cells dynamically change their transcriptional state over time as multiple gene programs - coordinated sets of genes that work together to perform specific biological tasks - are activated and repressed through a complex network of interactions. Dynamic changes in transcriptional cell states underlie fundamental biological processes, such as embryo development, tissue regeneration, and the onset and progression of cancer. A common approach to characterize cell state dynamics is to perform RNA sequencing at multiple time points, either at the ‘bulk’ or single-cell level, resulting in a series of gene expression profiles that represent the underlying cell states and that typically form continuous trajectories in latent space. However, the resulting expression profiles are typically a combination of multiple time-dependent components, where each component is associated with distinct biological processes and gene programs. A major challenge is to “deconvolve” these temporal components. This involves characterizing each component, identifying its associated gene programs, and inferring its relative contribution to the overall gene expression profiles at each time point.

Here, we demonstrate the use of unsupervised machine learning to reveal the temporal components in a simple model biological system. We first downloaded a dataset of bulk RNAseq measurements performed by Dominguez *et al*. on synchronized HeLa cells at 14 consecutive time points - approximately two cell cycles - following release from cell cycle arrest (Dominguez, Tsai, Gomez, et al. 2016; Dominguez, Tsai, Weatheritt, et al. 2016). Then, we performed Fourier analysis and found that most genes have a periodicity of either one or two cycles over time. Next, we used topic modeling, an unsupervised machine learning technique, and found that the series of gene expression profiles can be represented as a mixture of three components, two of which are periodic over time and correspond to the G1-S and G2-M phases of the cell cycle, and a third topic with a transient temporal behavior, that is associated with the immediate-early response gene program, regulation of cell proliferation, and cervical cancer. This study demonstrates the potential of machine learning algorithms to deconvolve hidden temporal components in complex biological systems, with potential applications for early disease detection, as well as monitoring disease progression and recovery processes.

## MATERIALS AND METHODS

### Datasets and Preprocessing

The RNAseq dataset generated by Dominguez *et al*. (Dominguez, Tsai, Gomez, et al. 2016; Dominguez, Tsai, Weatheritt, et al. 2016) was downloaded from GEO (accession number GSE81485) using SRA tools. STAR (Dobin et al. 2013) was used to align the reads to the human reference genome (hg38) and to obtain the raw counts matrix. DESeq2 (Love, Huber, and Anders 2014) was used to obtain the normalized counts matrix. Genes whose sum of counts across all 14 samples was below 10 were removed.

### RNA velocity

Velocyto (La Manno et al. 2018) was used to obtain the read counts of the spliced and unspliced mRNA for each sample. DESeq2 normalization was performed for the spliced and unspliced counts matrixes combined.

### Fourier Transform

For each gene, a Fourier analysis was performed using the R function “periodogram()” from the “TSA” (Time Series Analysis) R package. This function returns a vector of frequencies and a vector of scores (spectral densities) for each frequency. For each gene we found the “dominant frequency”, that is, the frequency with the highest score, and calculated a “dominant frequency score” which measures the degree to which the score of this frequency is higher than the scores of the other frequencies. Specifically, if for a particular gene the highest score is S1 and its matching frequency is F1, the second highest score is S2 and its frequency is F2, the third ranking score is S3 and its frequency is F3 etc., then the “dominant frequency” for that gene is F1, and the “dominant frequency score” is: *S*1/(*S*2 + *S*3). The significance of the dominant frequency scores was tested by comparing them to scores derived from randomized datasets, which were obtained by randomly shuffling the order of counts for each gene.

### Topic modeling

Topic modelling was performed on the normalized counts matrix using the R functions fit_topic_model(), diff_count_analysis(), and structure_plot() from the “fastTopics” R package (Carbonetto et al. 2021). We fitted a topic model with k=3 topics that we labeled as “k1”, “k2”, and “k3”. Using these functions, we calculated the three probability distributions *p*(*gene*|*k*1), *p*(*gene*|*k*2), and *p*(*gene*|*k*3) for every gene in each one of the three topics, as well as the probabilities *p*(*k*1|*sample*), *p*(*k*2|*sample*), and *p*(*k*3|*sample*) for each topic in every one of the fourteen samples. We also tested higher numbers of topics but the results did not change significantly.

One way to visualize the association between a specific gene and each of the three topics is to calculate the “posterior probabilities” *p*(*k*1|*gene*), *p*(*k*2|*gene*), and *p*(*k*3|*gene*) and compare them. In order to do this, we used Bayes’ theorem:

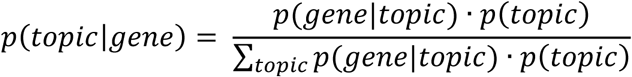

where, for simplicity, we set the prior *p*(*topic*) = 1/3 for each topic.

### Gene Ontology (GO) enrichment analysis

GO enrichment analysis was performed with Toppgene (Chen et al. 2007) (https://toppgene.cchmc.org/). Gene set enrichment analysis was performed with GSEA (Subramanian et al. 2005) (see Supplementary data for run details).

## RESULTS

### RNA velocity analysis of periodically expressed genes reveals a time lag between spliced and un-spliced mRNA

We downloaded a published dataset of RNAseq measurements that were performed by Dominguez *et al*. (Dominguez, Tsai, Gomez, et al. 2016; Dominguez, Tsai, Weatheritt, et al. 2016). In that study, HeLa cells were first synchronized by a double thymidine block. Then, following release from cell cycle arrest, RNA was collected and sequenced at 14 consecutive time points which corresponds to approximately two cell cycles (Figure 1A, Table S1). The authors also identified a set of 67 “core” cell cycle related genes with periodic behavior, and categorized them as “G1-S related” or “G2-M related” according to the stage at which their expression is maximal (Table S1).

**Figure 1:**
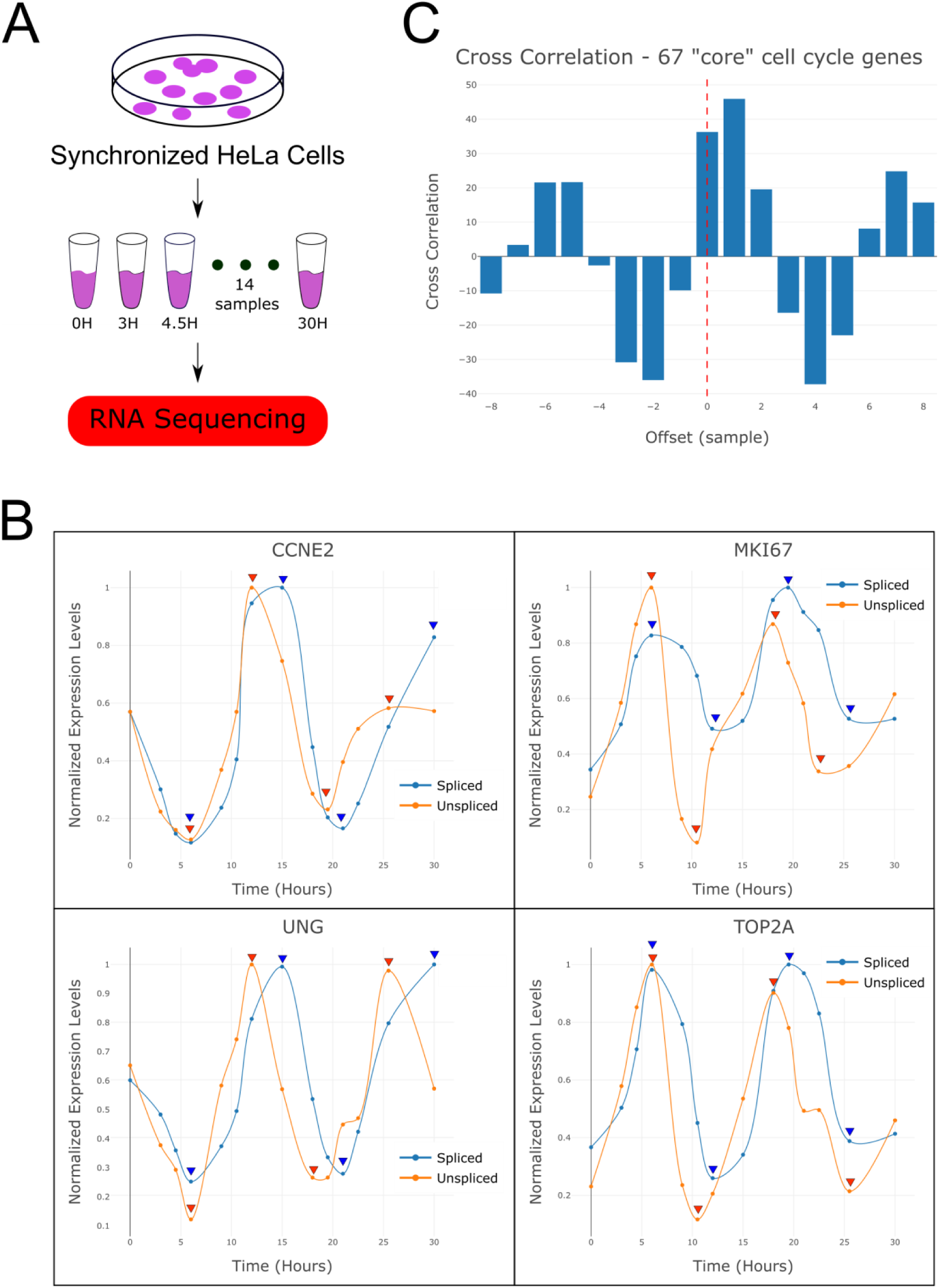
RNA velocity analysis of periodically expressed genes reveals a time lag between spliced and un-spliced mRNA. (A) Shown is a sketch of the experiment performed by Dominguez *et al*. from which the RNAseq dataset was produced. HeLa cells were sampled at 14 consecutive time points following release from cell cycle arrest and their RNA was sequenced. (B) Shown are expression levels over time of spliced and unspliced mRNA for selected genes that are known to be over-expressed at the G1-S phases (CCNE2, UNG) or the G2-M phases (MKI67, TOP2A) of the cell cycle. These indicate that the cells completed approximately two cell divisions during the experiment. It can be seen that the yet-unspliced mRNA again precedes the spliced mRNA, as expected. Expression levels were normalized to their maximum and spline curves were plotted through the data points to assist visibility. (C) A cross-correlation plot between the spliced and un-spliced mRNA levels averaged over the 67 “core” cell cycle genes (i.e. periodically expressed genes identified by Dominguez *et al*.) shows a significant time lag between the spliced and yet un-spliced mRNA.

We started by manually inspecting the expression levels of spliced and un-spliced mRNA of “core” cell cycle genes such as CCNE2 and UNG (G1-S related) and MKI67 and TOP2A (G2-M related, Figure 1B) (La Manno et al. 2018). As expected, we observed that the yet-unspliced mRNA typically precedes the spliced mRNA by a single sampling interval, which corresponds to a time interval of approximately 1.5-3 hours. We confirmed this by performing a cross-correlation between the spliced and un-spliced mRNA levels averaged over the 67 “core” cell cycle genes (Figure 1C).

### Fourier analysis identifies sets of genes with potentially transient and oscillatory behaviors over time

To test the periodicity of every gene in the transcriptome, we performed Fourier analysis. For each gene we plotted its “dominant frequency”, that is, the frequency with the highest spectral density, versus its “dominant frequency score”, which measures the degree to which the score of this frequency is higher than the scores of the other frequencies (see Methods and Figure 2A). We found that the first and second dominant frequencies - that correspond to either one or two whole cycles – contain genes with scores that are higher than those within other dominant frequencies (Figure 2A). Moreover, we observed that genes with dominant frequencies corresponding to three cycles and higher have scores comparable to those derived from randomized datasets, which were generated by randomly shuffling the order of counts for each gene (Figure S1). This suggests that most of the information in our dataset is contained in the expression levels of genes with a periodicity of either one or two cycles, indicating potential transient or oscillatory behavior, respectively.

**Figure 2:**
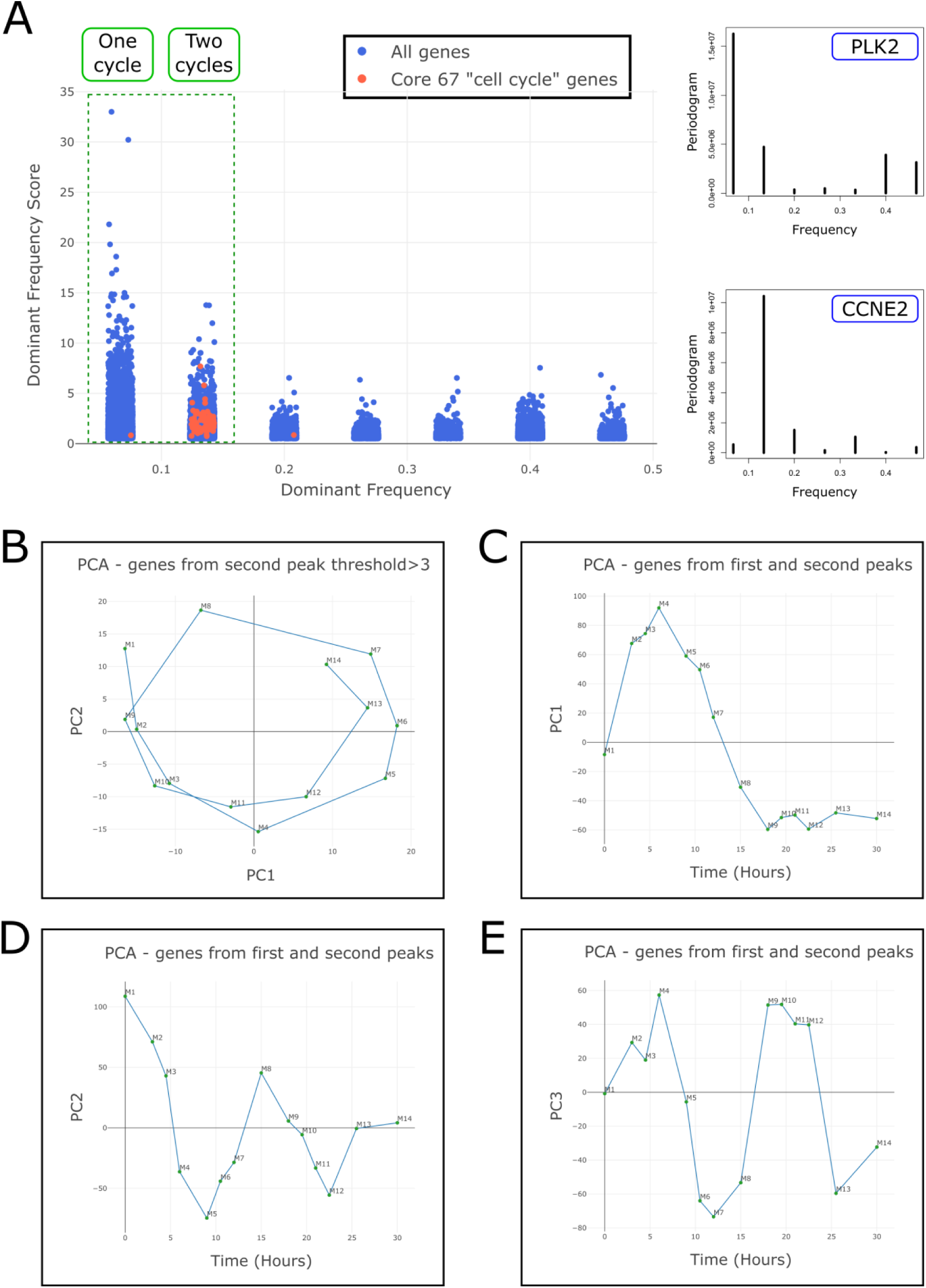
Fourier analysis identifies sets of genes with potentially transient and oscillatory behaviors over time. (A) For each gene, a periodogram was calculated (see specific examples for PLK2 and CCNE2 in the right panels) and its “dominant frequency” and “dominant frequency score” were plotted (left panel, see Methods). It can be seen that almost all of the “core” cell cycle genes (65 out of 67 red dots) have a dominant frequency corresponding to two whole cycles. Likewise, the first and second dominant frequencies contain genes with scores that are higher than those within other dominant frequencies, whereas genes in the 3rd dominant frequency and onwards have scores that are similar to those of randomly shuffled datasets (Figure S1). This indicates that the genes within the first and second dominant frequencies contain most of the periodic information in our dataset. (B) When we select genes from the second dominant frequency with scores > 3 and perform PCA, the samples form a circular trajectory in latent space that completes almost two cycles. (C-E) When we select genes from the first and second dominant frequencies and perform PCA, the samples form a trajectory in latent space with a transient behavior along PC1 axis and periodic behaviors along the PC2 and PC3 axes (that also create a circular trajectory in the PC2 vs. PC3 plane).

We next selected genes from the second dominant frequency (with scores > 3) and performed PCA. We found that the samples form a circular trajectory in latent space that corresponds to almost two complete cell cycles (Figure 2B). Moreover, separating the spliced and yet-unspliced mRNA expression profiles results in two circular trajectories in latent space with a phase difference between them (Figure S2). When we selected the genes from the first and second dominant frequencies we found that the samples form a trajectory in latent space with two patterns of behavior: a transient pattern along PC1 (Figure 2C) and an oscillatory pattern along PC2 and PC3 (Figures 2D-E).

### Topic modeling reveals three temporal components, two of which are periodic, corresponding to the G1-S and G2-M phases of the cell cycle, and a third, transient component, related to immediate-early response, regulation of cell proliferation, and cervical cancer

“Topic models” are used in computer science to describe collections of documents that contain varying proportions of words from a predetermined number k of different “topics” (Blei, Ng, and Jordan 2003; Dey, Hsiao, and Stephens 2017). By fitting a topic model to a set of documents, it is possible to discover both the k latent topics (i.e., the probability of occurrence of each word from the vocabulary within a topic) and their proportions in each individual document. In our case we assume that each cell state (gene expression profile) at a given time point is a weighted sum of k hidden components whose proportions vary with time. We therefore treat each cell state at a given time point as a document, each gene in the human genome as a word in the vocabulary, the presence of a specific RNA transcript within a cell state as the occurrence of a specific word in a document, the expression level (number of transcripts) of a gene in a cell state as the number of occurrences of a word in a document, and each latent component as a topic. Topic modeling can thus be used to find the k components (i.e. the probability of observing an RNA transcript from each gene within a component) and their proportions at each individual time point.

We fitted a topic model with k=3 topics to the 14 gene expression profiles and identified three components/topics with distinct temporal patterns, which we labeled as “k1”, “k2” and “k3” (Figure 3A and Table S2). We found that topics k1 and k3 have an oscillatory behavior over the course of both cycles following release from cell cycle arrest and are periodically over-expressed at specific phases of the cell cycle (Figures 3A-B). Topic k2 however is transiently over-expressed mainly during the first cycle following release from cell cycle arrest. We confirmed that the temporal pattern of each topic is also observed in selected genes associated with it. For example, the genes CCNE2 and UNG whose expression is periodic are associated with topic k1 (Figures 1B and 4A), whereas the genes MKI67 and TOP2A whose expression is also periodic, but with a different phase, are associated with topic k3 (Figures 1B and 4A). Likewise, the genes KIFC3, PHLDB2, and PLK2, which are associated with topic k2, are transiently over-expressed following release from cell cycle arrest (Figures 3C and 4A).

**Figure 3:**
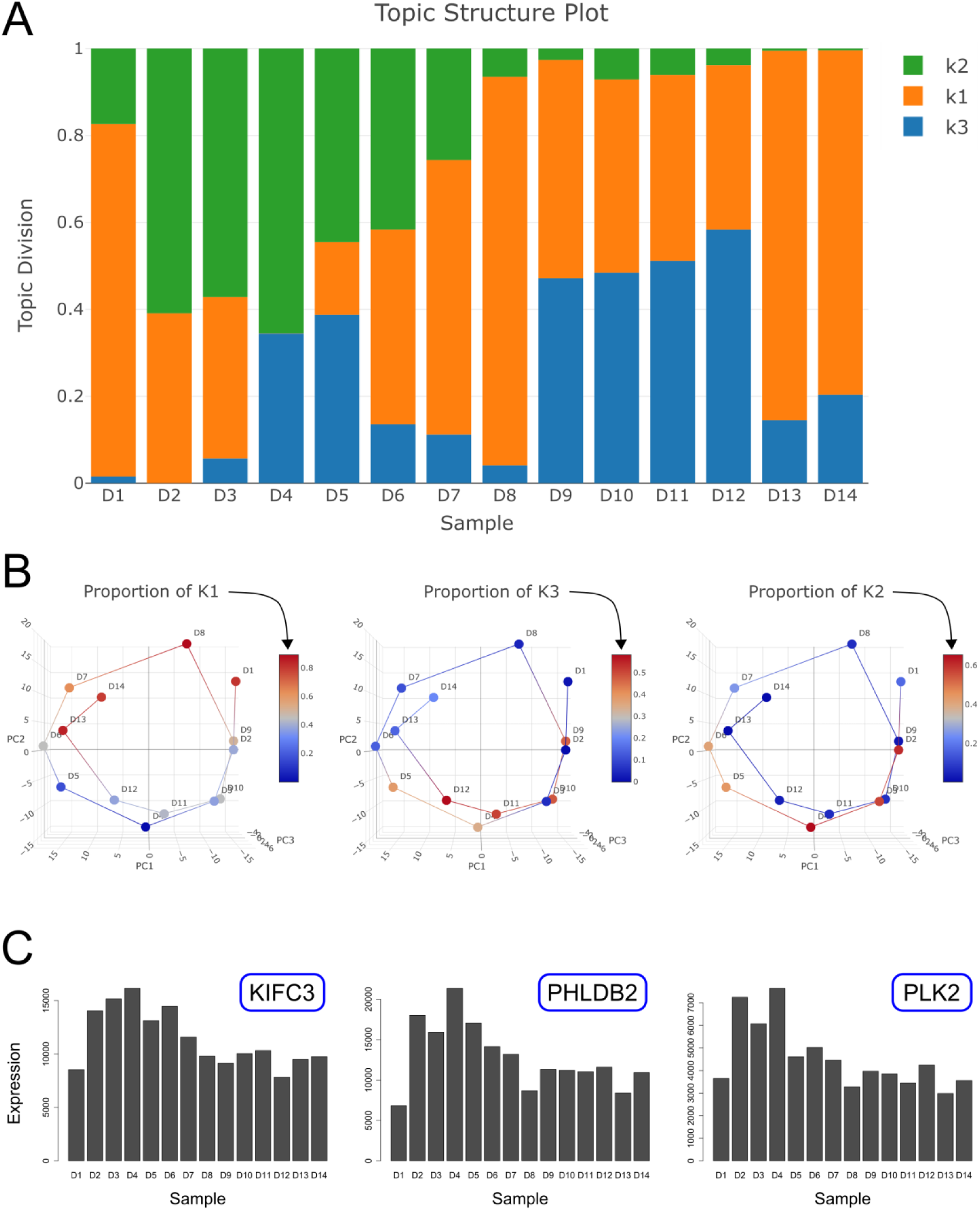
Topic modeling reveals three temporal components with periodic and transient behaviors over time. (A) Shown is a structure plot of the proportions of each component/topic within each sample. Topics k1 and k3 are periodically expressed over the course of both cycles (samples D1-D14), whereas topic k2 is transiently over-expressed mainly during the first cycle (samples D1-D7) following release from cell cycle arrest. (B) Shown is a visualization of the proportions of each topic along the circular trajectory in latent space representing the two cell cycles (red – high proportion, blue – low proportion). It can be seen that topics k1 and k3 are over-expressed in distinct phases during both cycles, whereas topic k2 is over-expressed during the first cycle only. Note that topic k1 is expressed mainly in the upper half of the circular trajectory, whereas topic k3 is expressed mainly in its lower half. (C) Shown are the expression levels of three representative genes that are associated with topic k2 and are transiently over-expressed following release from cell cycle arrest.

**Figure 4:**
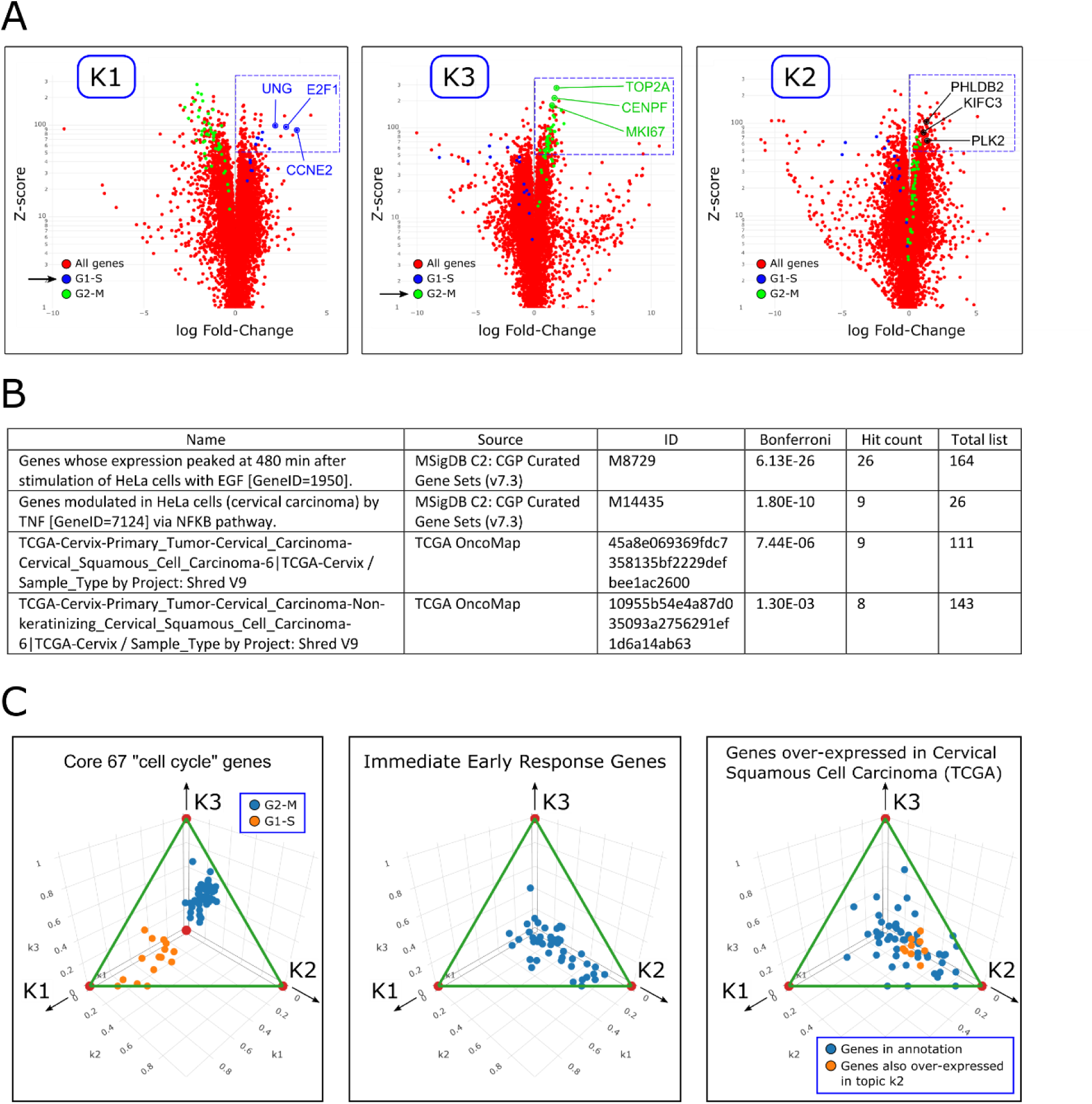
Gene Ontology (GO) enrichment shows that the two periodic components correspond to the G1-S and G2-M phases of the cell cycle, whereas the third transient component is related to immediate-early response, regulation of cell proliferation, and cervical cancer. (A) Shown are volcano plots illustrating the over-expression of each gene in each of the three topics. It can be seen that genes known to be over-expressed in the G1-S phases of the cell cycle are over-expressed in topic k1 (left panel), genes known to be over-expressed in the G2-M phases of the cell cycle are over-expressed in topic k3 (middle panel), and genes that are transiently over-expressed following release from cell cycle arrest are over-expressed in topic k2 (right panel). (B) GO enrichment analysis of genes that are over-expressed in topic k2 (Z-score>50) shows many of them are related to cervical cancer or involved in response to growth factors (Table S4). (C) Shown are the posterior probabilities *p*(*topic*|*gene*) for selected gene sets or specific GO terms. Each point represents a gene, and its components along the k1, k2, and k3 axes are the posterior probabilities *p*(*k*1|*gene*), *p*(*k*2|*gene*), and *p*(*k*3|*gene*), which represent the association between that gene and each of the three topics. It can be seen that topic k1 is highly associated with genes that are known to be over-expressed in the G1-S phases of the cell cycle (left panel) and that topic k3 is highly associated with genes that are known to be over-expressed in the G2-M phases of the cell cycle. Topic k2 is associated with genes that are related to the immediate-early response program and to cervical carcinoma (middle and right panels). Note that all points (=genes) are located on the triangle-shaped two-dimensional simplex since the sum of the posterior probabilities for each gene equals 1.

To identify and characterize the topics, we performed Gene Ontology (GO) enrichment analysis. For each topic, we selected genes that are significantly over-expressed and used them as input to ToppGene (Chen et al. 2009) (Figure 4A, Tables S3-S5). We found that topic k1 over-expresses genes related to the G1-S phase of the cells cycle (e.g. DNA replication and G1 to S cell cycle control) and that topic k3 over-expresses genes that are related to the G2-M phase of the cells cycle (e.g. mitosis and regulation of the G2/M transition). In contrast, topic k2 was found to over-express genes associated with the following gene programs: (i) immediate-early response (Herschman 1991; O’Donnell, Odrowaz, and Sharrocks 2012; Tullai et al. 2007; Winkles 1997) (Table S6 and Figure S3); (ii) regulation of cell proliferation (Table S4), and (iii) cervical carcinoma (Table S4).

We confirmed these observations by calculating the posterior probabilities p(k1|gene), p(k2|gene), and p(k3|gene) for genes belonging to specific gene sets and GO terms (Figure 4C). These probabilities represent the association between these genes and each of the three topics. Indeed, we observed that within the 67 “core” cell cycle genes, the G1-S related genes were associated with topic k1 and the G2-M related genes were associated with topic k3. Likewise, genes related to immediate-early response and cervical carcinoma were more strongly associated with topic k2.

## DISCUSSION

In this study we applied Fourier analysis and Topic modeling to a simple *in-vitro* model system in order to identify its latent components. We identified two components with a periodic temporal behavior (topics k1 and k3), that correspond to the G1-S and G2-M phases of the cell cycle, and a third component with a transient expression pattern (topic k2) that is associated with the immediate-early response gene program, regulation of cell proliferation, and cervical cancer.

Specific examples of genes that are transiently over-expressed following release from cell cycle arrest, and that have a high probability for being expressed in topic k2, are KIFC3, PHLDB2, and PLK2 (Figures 3C and 4A). These genes are known to be associated with cervical carcinoma (Table S4). Moreover, PLK2, a member of the polo-like kinase family, is also known to be associated with positive regulation of the cell cycle (de Cárcer, Manning, and Malumbres 2011) and is regarded as an immediate-early response gene (Winkles 1997) (Figure S17). Other examples include the genes JUN (C-Jun), FOS (c-Fos), FOSB, and FOSL1 (FRA-1), that are also transiently over-expressed following release from cell cycle arrest (Figure S13). The protein products of these genes are components of the AP-1 transcription factor complex that is thought to be involved in cell proliferation and cancer progression (Casalino et al. 2022; Milde-Langosch 2005). Furthermore, the genes JUN, FOS, and FOSB are also members of the immediate-early response gene family. The genes RELB, NFKB1, and NFKB2, are also transiently over-expressed following release from cell cycle arrest (Figure S18). These genes are members of the NF-kB family of transcription factors and are known to induce target genes involved in initiation and progression of cancer (Costa et al. 2016; Tilborghs et al. 2017). We also observed that the apoptosis inhibitors BIRC2 and BIRC3 (Silke and Vaux 2001) are transiently over-expressed following release from cell cycle arrest (Figure S27), whereas the gene BIRC5 (Survivin), which is a regulator of the mitotic cell cycle, shows a periodic pattern of expression. Additional examples are shown in Figures S4-S27.

The association of immediate-early response genes with topic k2 is not very surprising, since these genes are known to be rapidly and transiently transcribed in response to stimulation of cells by serum or growth factors such as EGF, subsequently promoting phenotypic changes such as proliferation, differentiation, and survival (Winkles 1997). In our study, it is likely that these genes facilitate cell-cycle reentry, promoting the transition from the quiescent G0 phase into the G1 phase. The association of topic k2 with cervical cancer is less straightforward. One possible explanation for this association is that, following release from cell cycle arrest, the cells enter a transient state lasting approximately twelve hours (samples D1-D7), during which they activate molecular mechanisms to resume the cell cycle. In normal tissue differentiation, cell cycle exit and lineage-specification are often regulated by tissue-specific mechanisms (Theilgaard-Mönch et al. 2022). It is plausible that cell cycle re-entry is similarly regulated by mechanisms specific to HeLa cells and cervical cancer, the tissue from which they originate.

For some of the 67 “core” cell cycle genes we observed that the spliced and unspliced mRNA levels arrive at their first peak (either maximum or minimum) at the same time (Figure 1B), and only after a few hours the typical time lag between the spliced and un-spliced mRNA expression is acquired. This indicates that initially, during the few hours after release from cell cycle arrest, the spliced and un-spliced transcripts are synchronized in these genes. Additional evidence for this gradual acquisition of time lag between spliced and un-spliced mRNA can be seen for larger sets of genes with periodic behavior (Figure S2A-B). This indicates that in the transient cell state following release from cell cycle arrest, the transcriptional timing mechanisms operate in a way that is different from the normal cycling state. One possible explanation is that during this period, the primary mRNA transcribed from these genes is spliced, processed, and translated with minimal delay in order to rebuild the necessary mechanisms for cell cycle re-entry as soon as possible. According to this hypothesis, splicing mechanisms participate in the immediate-early response mechanism. Further studies at higher time resolution are required to systematically test this hypothesis and inspect the relationship between mRNA splicing and the immediate-early response gene program.

## ACKNOWLEDGEMENTS

We wish to thank Shahar Alon and all the member of our lab for useful comments and suggestions.

## DECLARATION OF INTEREST STATEMENT

The authors have declared that no competing interests exist.

## AUTHOR CONTRIBUTIONS

Study initiation and conception – T.M. and T.K.; Data analysis - T.M. and T.K.; Topic modeling – T.M, J.G., and T.K.; Other intellectual contribution – Y.T.; Manuscript writing – T.M. and T.K.

## FUNDING

T.M., Y.T., and T.K. were supported by the Israel Science Foundation (ICORE no. 1902/12 and Grants no. 1634/13, 2017/13, and 1814/20), the Israel Ministry of Health (Grant no. 3-10146), the EU-FP7 (Marie Curie International Reintegration Grant no. 618592), the Data Science Institute at Bar-Ilan University (seed grant), the ICRF (Grant no. 19-101-PG), the Israel Ministry of Science (Grant no. 3-16220), the Israel Ministry of Justice (Veadat Haezvonot), and the Israel Cancer Association (Grant no. 20240114). The funders had no role in study design, data collection and analysis, decision to publish, or preparation of the manuscript.

## APPENDICES

**Supplementary information:** Supplementary text and figures.

**Table S1:** The gene expression dataset from Dominguez *et al*., including the sampling time points and the list of 67 “core” cell cycle genes.

**Table S2:** The three topics discovered and their proportions at each time point. **Table S3:** GO enrichment analysis for genes that are over-expressed in topic k1. **Table S4:** GO enrichment analysis for genes that are over-expressed in topic k2. **Table S5:** GO enrichment analysis for genes that are over-expressed in topic k3. **Table S6:** A list of immediate-early response genes from Tullai *et al*.

**Program:** A compressed directory containing programs and datasets for data visualization.

## Notes

### Competing Interest Statement

The authors have declared no competing interest.

### Summary of Updates

1. Additional results and biological insight: We found that the transient temporal component ("topic k2") is also associated with the "immediate-early response" gene program. We performed additional statistical analysis in the "Results" section. 2. Changes of terminology in the "Background" section: RNA sequencing measurements performed at multiple time points results in a series of gene expression profiles that represent the underlying "cell states". The expression profiles are a combination of multiple time-dependent "components", where each component is associated with distinct biological processes and gene programs. 3. In the discussion section we review the literature on selected genes associated with the immediate-early response program and with cervical carcinoma.

