## Supplementary text and figures. for "Unsupervised machine learning reveals temporal components of gene expression in HeLa cells following release from cell cycle arrest"

[2]Correspondence to:

Tomer Kalisky

**FIGURES**

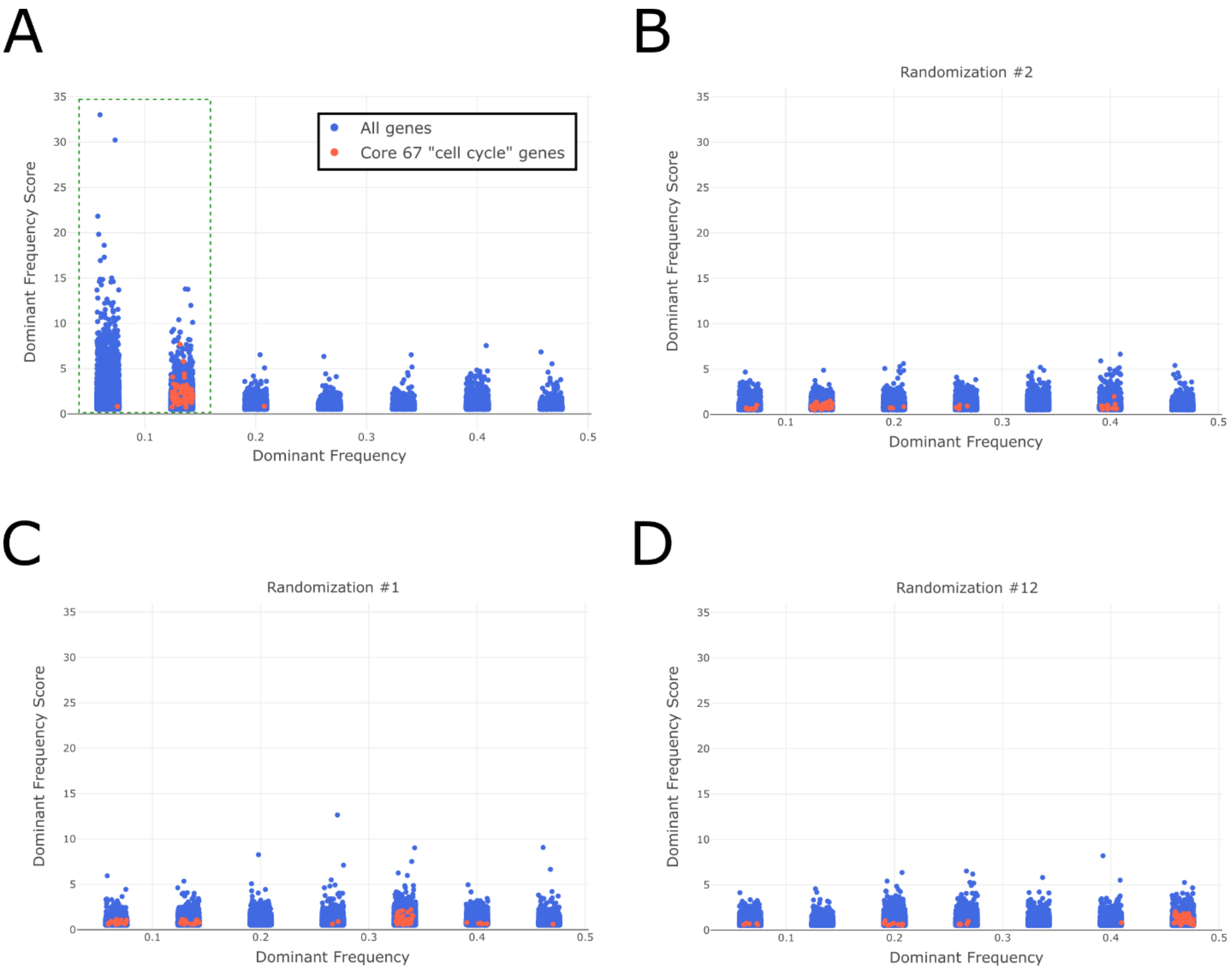


**Figure S1: Fourier analysis identifies sets of genes with potentially transient and oscillatory behaviors over time.**

(A) A periodogram was calculated for each gene, and its dominant frequency vs. dominant frequency score were plotted (see Methods). It can be seen that genes (=dots) within the first and second dominant frequencies have scores that are higher than genes within other dominant frequencies. Likewise, the genes within the 3rd dominant frequency and onwards have scores that are similar to those derived from randomized datasets, which were generated by randomly shuffling the order of counts for each gene (B-D). This indicates that the genes within the first and second dominant frequencies contain most of the periodic information in our dataset.

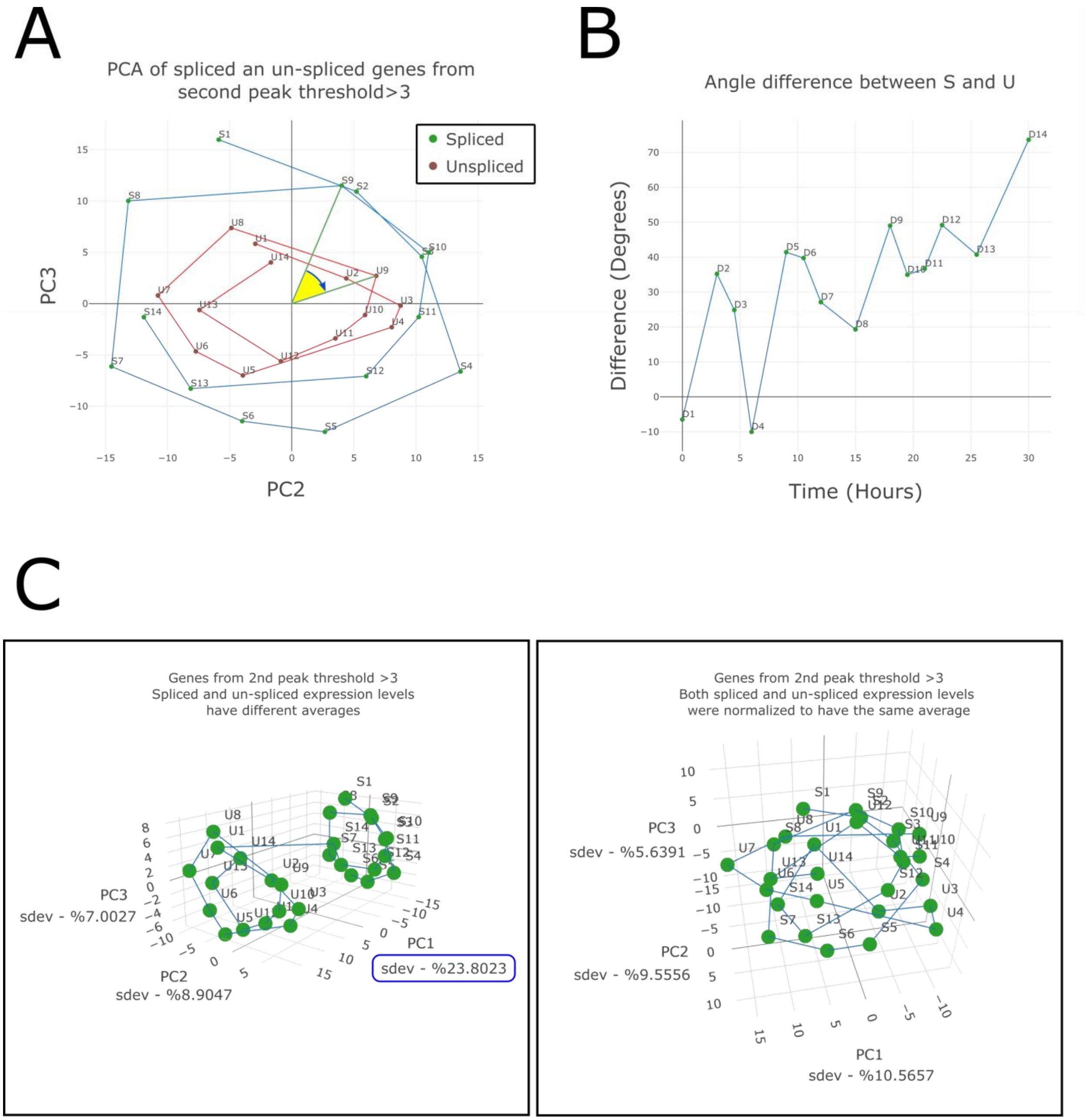


**Figure S2: RNA velocity analysis of periodically expressed genes reveals a time lag between spliced and un-spliced mRNA.**

(A) After performing Fourier analysis, selecting only genes from the second dominant frequency with scores > 3, and performing PCA, the samples form a circular trajectory in latent space that represents almost two complete cell cycles. Further separation into spliced (S1, S2, S3, …) and un-spliced (U1, U2, U3, …) mRNA expression profiles results in two circular patterns in latent space with a rotation angle difference between them (note: the unspliced circle was artificially made smaller in order to assist visibility) (B) This difference in rotation angle appears to start from zero, increase, and then stabilize over time at some constant value. However, longer and more frequent measurements will be needed to validate this observation. (C) Note that the two circles representing the spliced and un-spliced data points are oriented in parallel to the PC2 vs. PC3 plane and are widely separated along the PC1 axis as a result the difference in average expression between the spliced and un-spliced mRNA in each gene (left panel). Normalizing the spliced and un-spliced expression profiles to have the same average expression in each gene results in both circles collapsing on the same plane in latent space, that is, the PC1 vs. PC2 plane, with negligible separation along the PC3 axis (right panel).

A

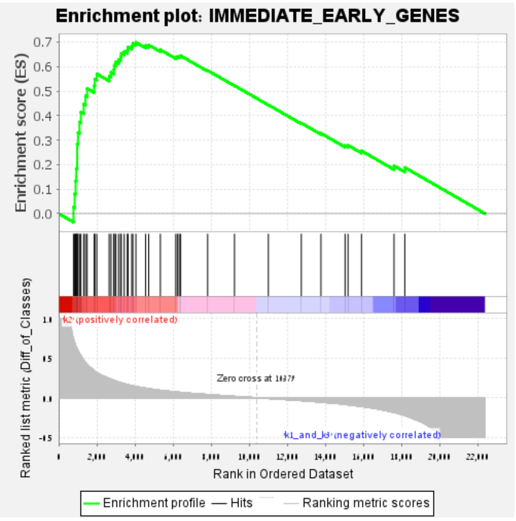


| | |
|---|---|
| Dataset | 2024.06.11 Table S2 - The three topics and their proportions.2024.06.k2_vs_k1_and_k3.cls #k2_versus_k1_and_k3.k2_vs_k1_and_k3.cls #k2_versus_k1_and_k3_repos |
| Phenotype | k2_vs_k1_and_k3.cls#k2_versus_k1_and_k3_repos |
| Upregulated in class | k2 |
| GeneSet | IMMEDIATE_EARLY_GENES |
| Enrichment Score (ES) | 0.69883245 |
| Normalized Enrichment Score (NES) | 2.1219182 |
| Nominal p-value | 0.0 |
| FDR q-value | 0.0 |
| FWER p-Value | 0.0 |

B

C

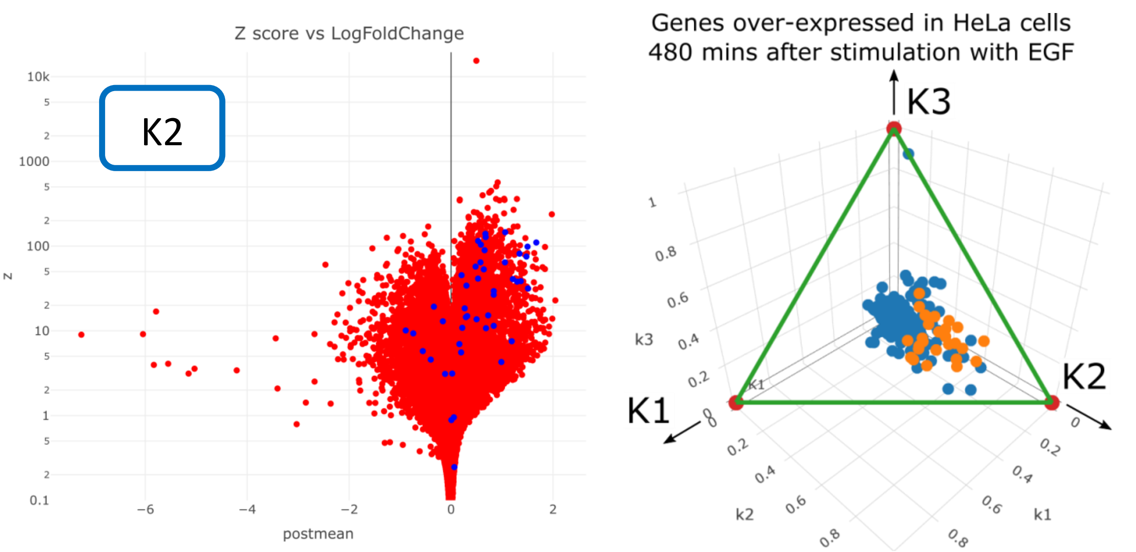


**Figure S3: Immediate-early genes are associated with topic k2.**

(A) A GSEA enrichment plot shows that the immediate-early genes are significantly over-expressed in topic k2 vs. topics k1 and k3 (Enrichment Score = 0.69, Normalized Enrichment Score = 2.12, p_value < $10^{-3}$, see supplementary data). (B) A volcano plot shows that a large proportion of immediate-early genes are over-expressed in topic k2. Each data point represents a gene, where the immediate-early genes are colored in blue. (C) A posterior probability plot demonstrates similar enrichment for a set of genes that showed transient over-expression following stimulation with EGF (epidermal growth factor), a component of calf serum that also stimulates expression of immediate-early genes (Winkles 1997).

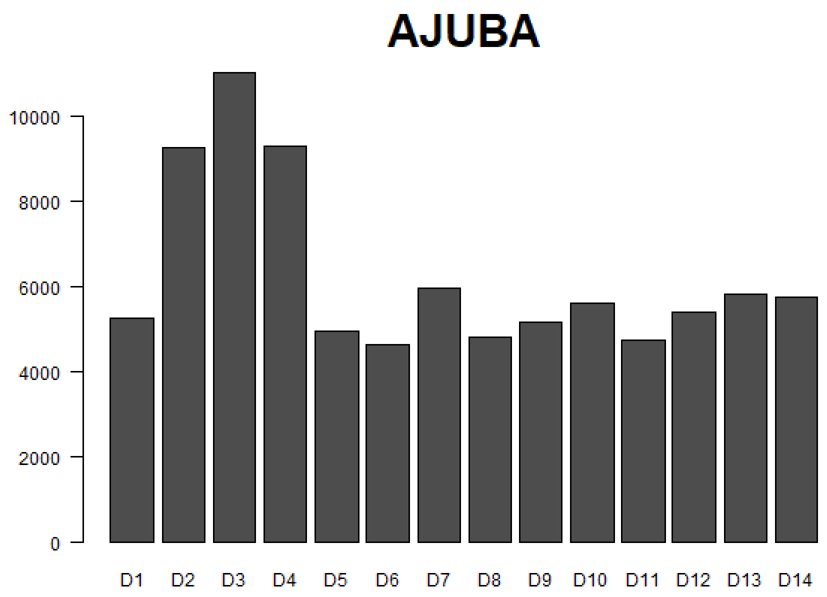


**Figure S4: The gene AJUBA is transiently over-expressed following release from cell cycle arrest and is associated with topic k2.**

The gene AJUBA was previously found to be over-expressed in cervical cancer (Bi et al. 2018) and its depletion was found to cause S-phase delay in cell lines (Kalan, Matveyenko, and Loayza 2013).

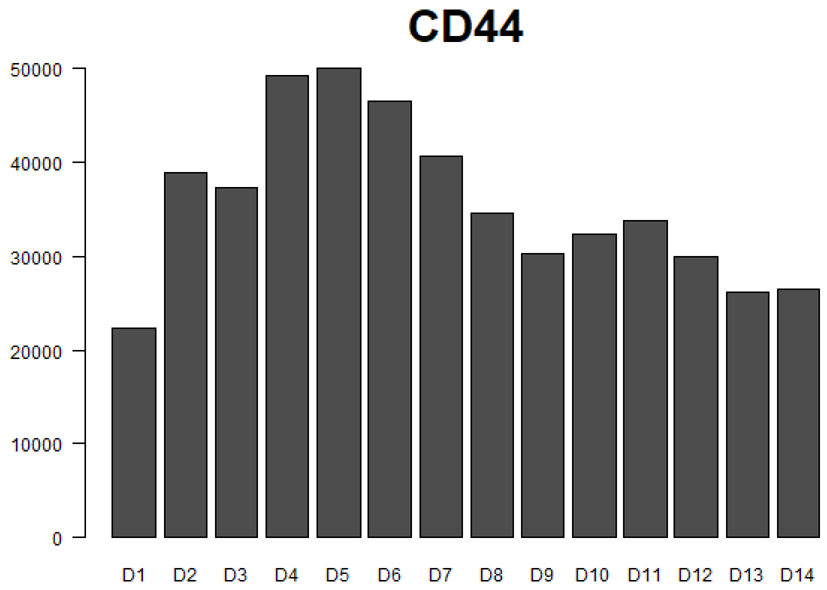


**Figure S5: The gene CD44 is transiently over-expressed following release from cell cycle arrest and is associated with topic k2.**

The expression of the gene CD44 was previously found to increase with progression from normal cervical epithelium to high-grade squamous intraepithelial lesions, and further with progression to squamous cell carcinoma of the cervix (Mehdi, Raju, and Sheela 2023).

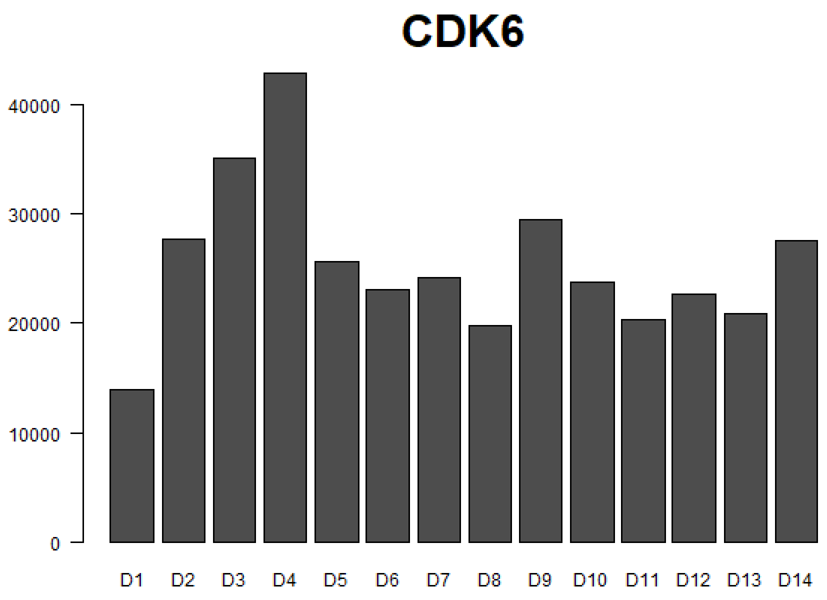


**Figure S6: The gene CDK6 is transiently over-expressed following release from cell cycle arrest and is associated with topic k2.**

The gene CDK6 is thought to be associated with the onset of cell cycle progression (Meyerson and Harlow 1994). CDK6 was previously found to be significantly over-expressed in cervical tumors, relative to both normal cervical epithelium and cervical intraepithelial neoplasia (CIN) (Huang et al. 2021).

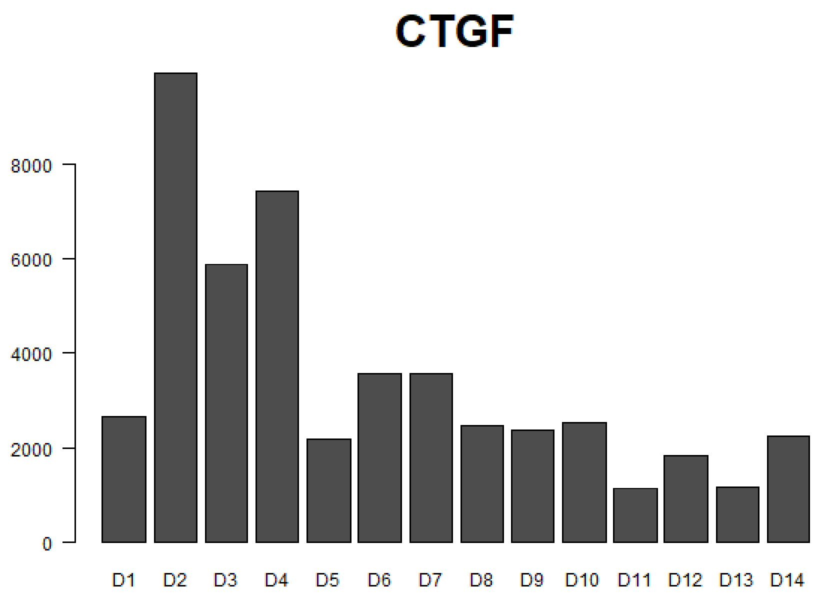


**Figure S7: The gene CTGF (CCN2) is transiently over-expressed following release from cell cycle arrest and is associated with topic k2.**

The gene CTGF (CCN2) is an immediate-early response gene (Tullai et al. 2007). This gene was previously found to be dysregulated in cervical cancer, showing downregulation in cancer compared to normal tissues (Xie et al. 2012), and upregulation in late stage cancer compared to early stage cancer (Wong et al. 2006).

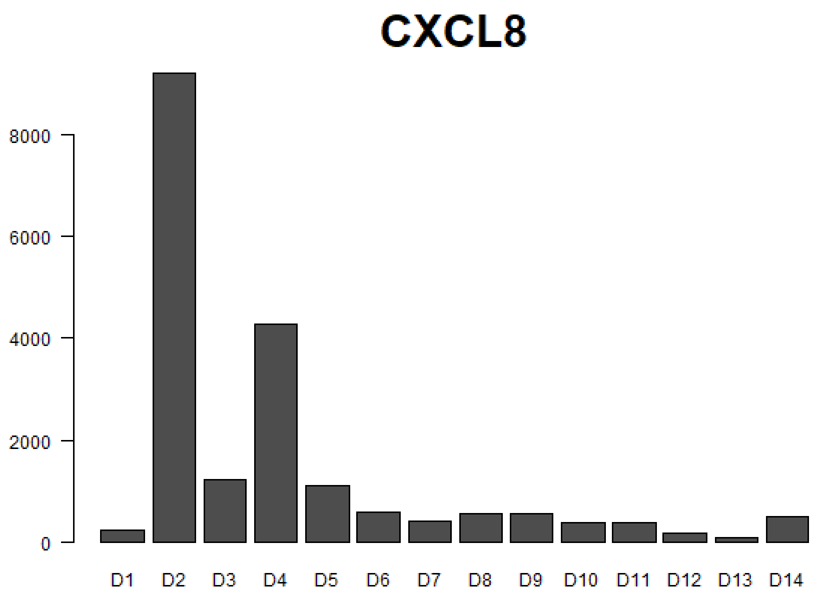


**Figure S8: The gene CXCL8 is transiently over-expressed following release from cell cycle arrest and is associated with topic k2.**

The gene CXCL8 was previously found to be overexpressed in cervical cancer biopsies relative to normal tissues (Fernandez-Avila et al. 2023), and higher expression levels of CXCL8 were found to be related to a worse prognostic survival.

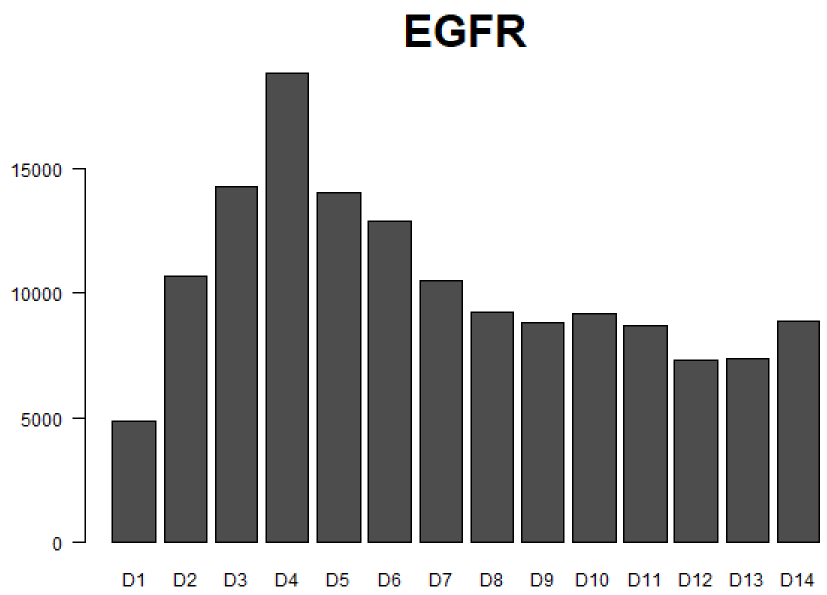


**Figure S9: The gene EGFR is transiently over-expressed following release from cell cycle arrest and is associated with topic k2.**

The gene EGFR, a receptor tyrosine kinase that converts extracellular cues into cellular responses such as cell proliferation, was previously found to be significantly over-expressed in patients with invasive cervical cancer, in both the primary tumors and the lymph node metastases (J. W. Kim et al. 1996; Shen et al. 2008).

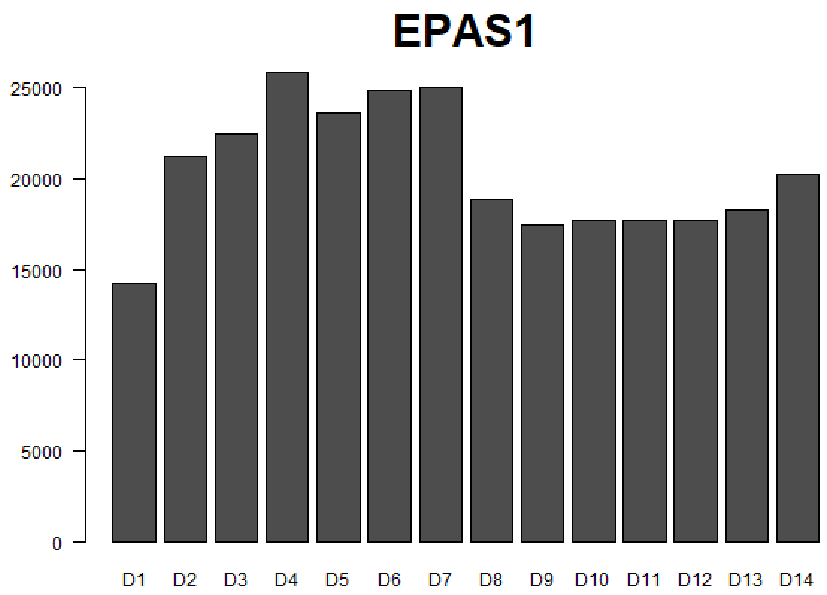


**Figure S10: The gene EPAS1 (HIF2A) is transiently over-expressed following release from cell cycle arrest and is associated with topic k2.**

The gene EPAS1 (HIF2A), a transcription factor that regulates genes involved in response to low oxygen concentration, was previously found to be over-expressed in Cervical Squamous Cell Carcinoma with respect to normal cervical tissues (L. Zhang et al. 2016).

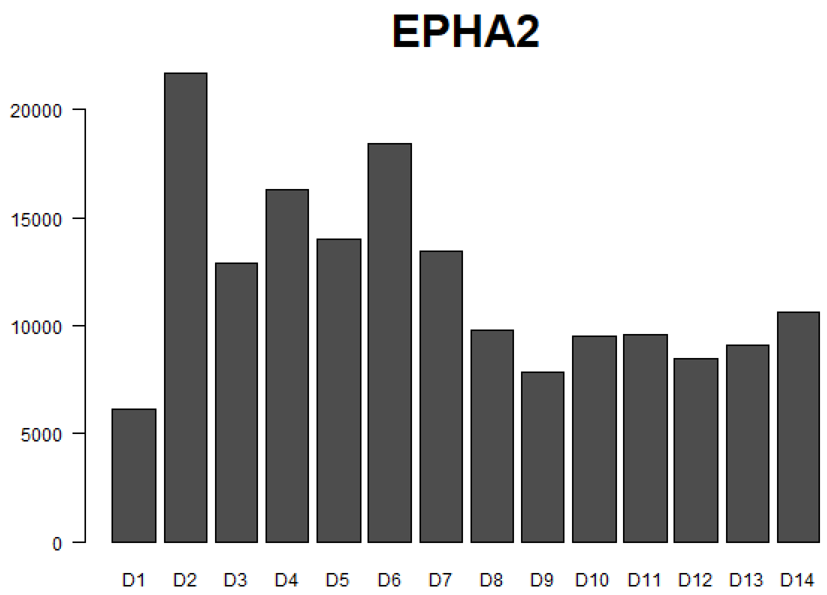


**Figure S11: The gene EPHA2 is transiently over-expressed following release from cell cycle arrest and is associated with topic k2.**

The gene EPHA2 was previously found to be highly expressed in cervical cancer relative to both normal cervical epithelium and cervical intraepithelial neoplasia (CIN) (Huang et al. 2021). Elevated levels of EPHA2 were also observed in advanced-stage tumors and tumors with lymph node metastasis.

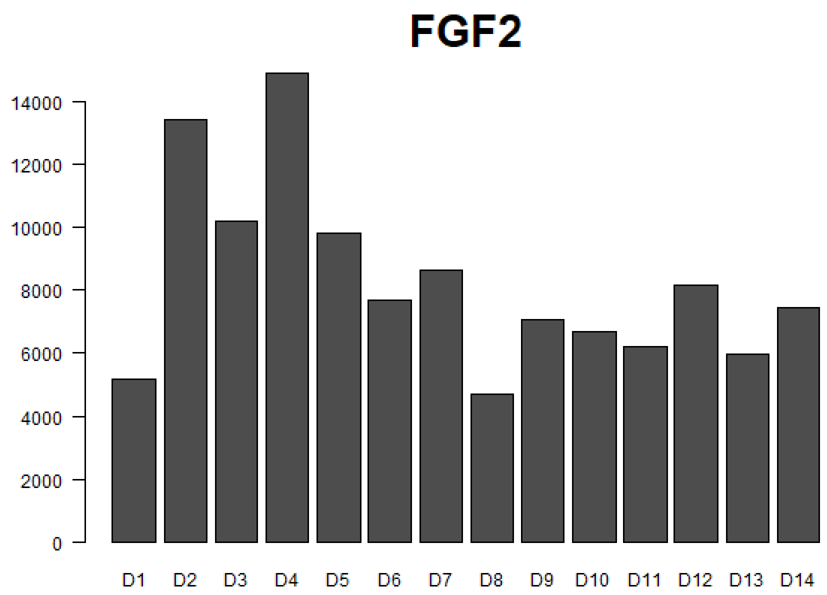


**Figure S12: The gene FGF2 is transiently over-expressed following release from cell cycle arrest and is associated with topic k2.**

The gene FGF2 (also known as basic fibroblast growth factor, or BFGF) was previously found to be highly expressed in advanced-stage cervical cancer, regardless of histological type (Fujimoto et al. 1997).

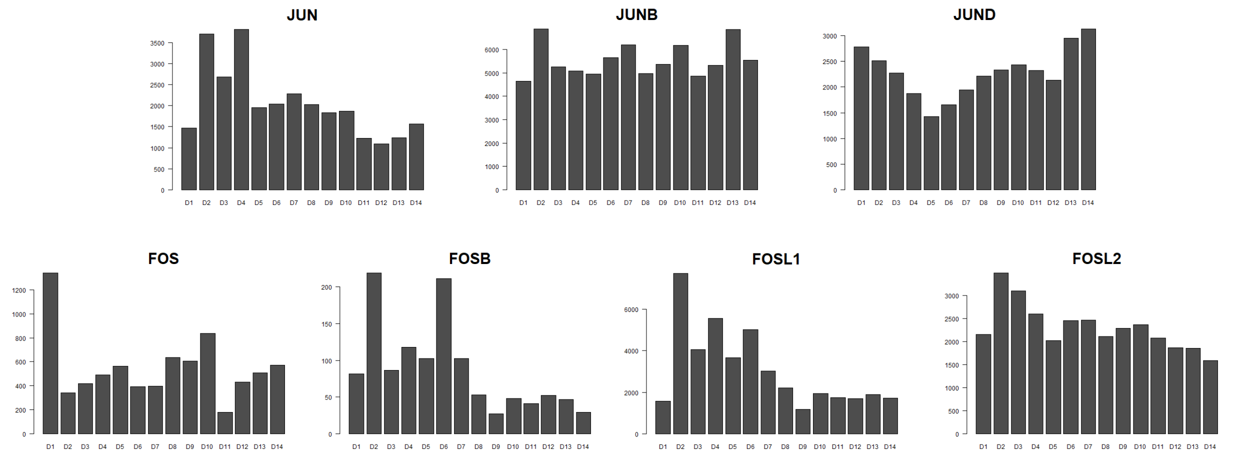


**Figure S13: The genes JUN (C-Jun) and FOSL1 (FRA-1) are transiently over-expressed following release from cell cycle arrest and are associated with topic k2.**

The proteins encoded by the genes JUN (c-Jun), JUNB, JUND, FOS (c-Fos), FOSB, FOSL1 (FRA-1), and FOSL2 (Fra-2) can heterodimerize to form the AP-1 complex, a transcription factor known to be involved in cell proliferation and cancer progression (Casalino et al. 2022; Milde-Langosch 2005; O'Donnell, Odrowaz, and Sharrocks 2012). Likewise, the genes JUN, JUNB, FOS, and FOSB were previously classified as immediate-early response genes (Tullai et al. 2007). It was previously found that downregulation of JUN inhibits the proliferation and invasion potential of HeLa cells (Yee, De Souza, and Khachigian 2013). Note, however, that the association of FOSL1 to cervical cancer progression not straightforward. For example, FOSL1 was found to gradually decrease during progression of normal tissue to early precancerous lesions and then to invasive cervical tumors (Prusty and Das 2005). Moreover, the high levels of FOSL1 expression in the early precancerous cervical lesions were found to be associated with low overall AP-1 binding activity.

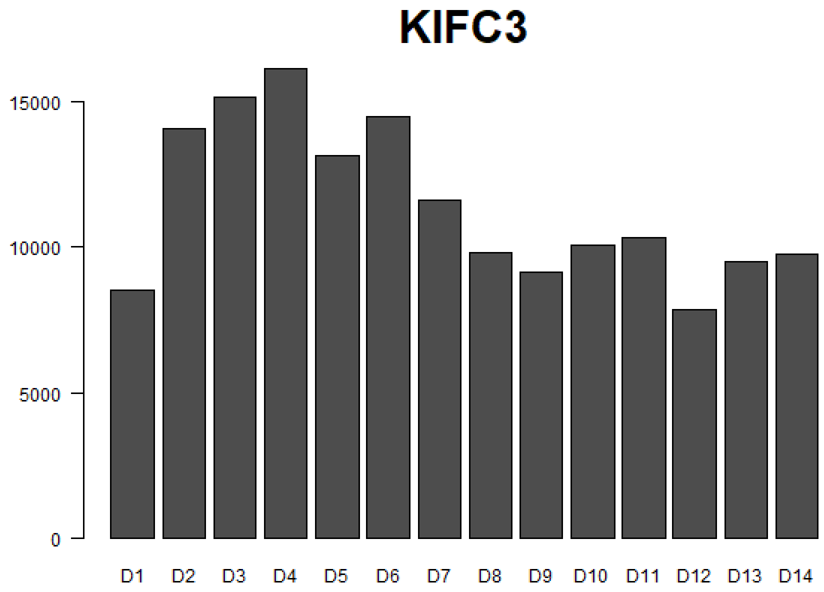


**Figure S14: The gene KIFC3 is transiently over-expressed following release from cell cycle arrest and is associated with topic k2.**

The gene KIFC3 was previously found to be associated with cell proliferation, migration, and invasion in colorectal cancer (Liao et al. 2022).

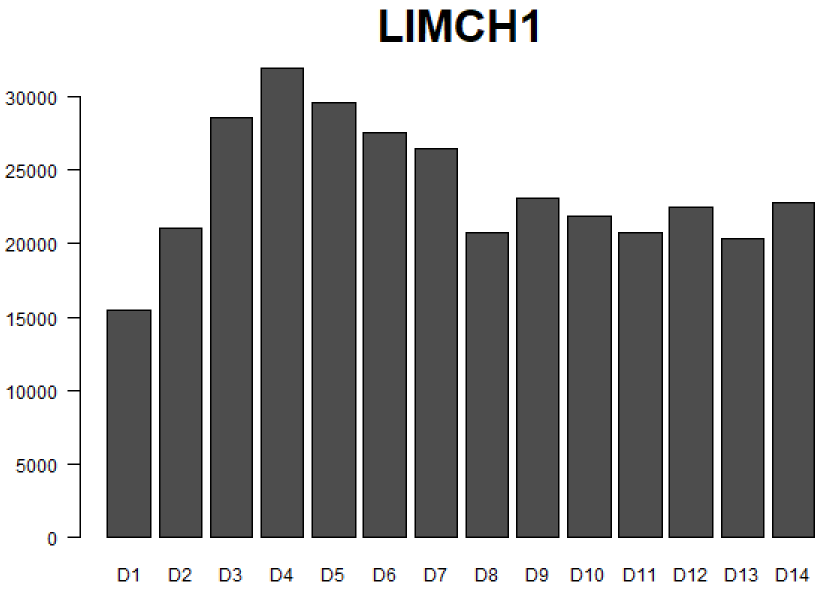


**Figure S15: The gene LIMCH1 is transiently over-expressed following release from cell cycle arrest and is associated with topic k2.**

High expression levels of the gene LIMCH1 were previously found to predict poor outcome in cervical cancer patients (Halle et al. 2021).

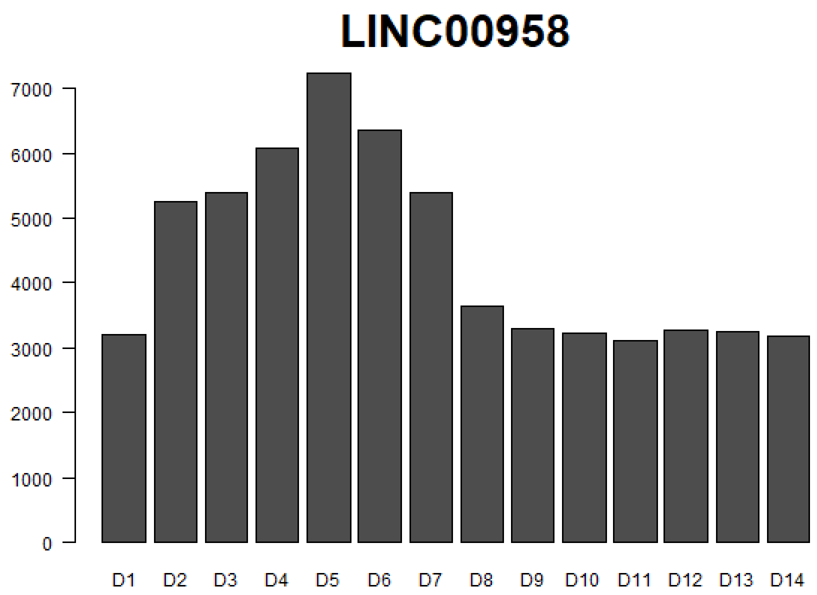


**Figure S16: The long noncoding RNA LINC00958 is transiently over-expressed following release from cell cycle arrest and is associated with topic k2.**

The long noncoding RNA LINC00958 was previously found to be overexpressed in cervical cancer and to be associated with cervical cancer cell proliferation and metastasis (L. Wang et al. 2020).

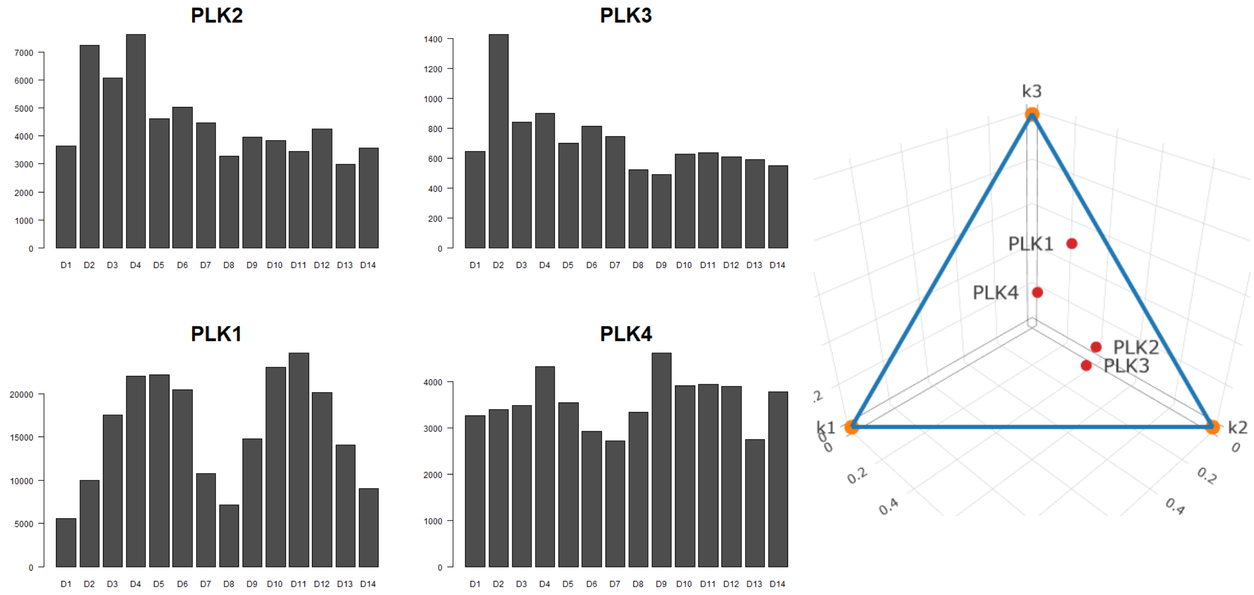


**Figure S17: The genes PLK2 and PLK3 are transiently over-expressed following release from cell cycle arrest and are associated with topic k2.**

The polo-like kinases PLK1, PLK2, PLK3, and PLK4, are associated with cell cycle regulation and progression (de Cárcer, Manning, and Malumbres 2011). PLK1, which is expressed in highly proliferating tissues, is known to perform multiple functions during the cell cycle (Weerdt and Medema 2006). On the other hand, PLK2 and PLK3, which have a more broad tissue distribution, are considered immediate-early response genes (Winkles 1997). In the dataset that we studied we observed that PLK2 and PLK3 are transiently over-expressed following release from cell cycle arrest, while PLK1 (and PLK4 to some extent) are modulated according to the cell cycle phase. The diagram of posterior probabilities (right) illustrates the association of PLK2 and PLK3 with topic k2, which represents transient expression following release from cell cycle arrest, and of PLK1 with topic k3, linked to the G2-M phases of the cell cycle.

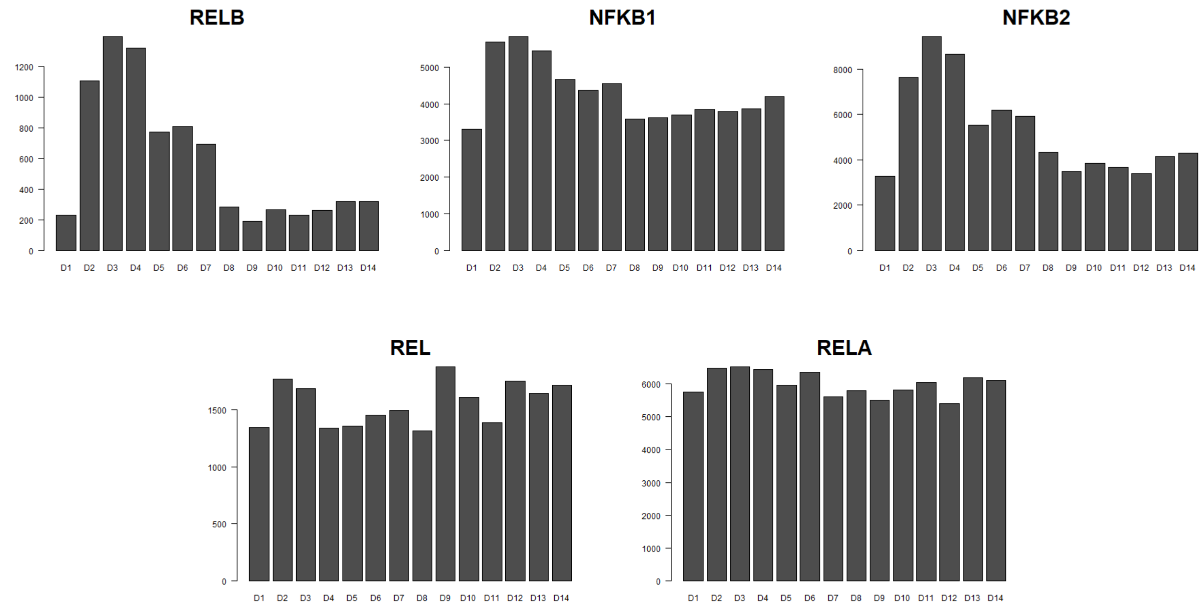


**Figure S18: The genes RELB, NFKB1, and NFKB2 are transiently over-expressed following release from cell cycle arrest and are associated with topic k2.**

The genes REL (c-Rel), RELA (p65), RELB, NFKB1 (p105/p50), and NFKB2 (p100/p52), are members of the NF-kB family of transcription factors that are known to form various homo- or heterodimers. Upon activation these transcription factors can translocate to the nucleus, bind to DNA cis-regulatory elements at enhancers and promoters, and induce target genes involved in initiation and progression of cancer (Costa et al. 2016; Tilborghs et al. 2017). In cervical cancer, increased expression of the nuclear (and thus active) forms of RELA (p65) and NFKB1 (p50) was found to be associated with tumor progression and metastasis (Li et al. 2009).

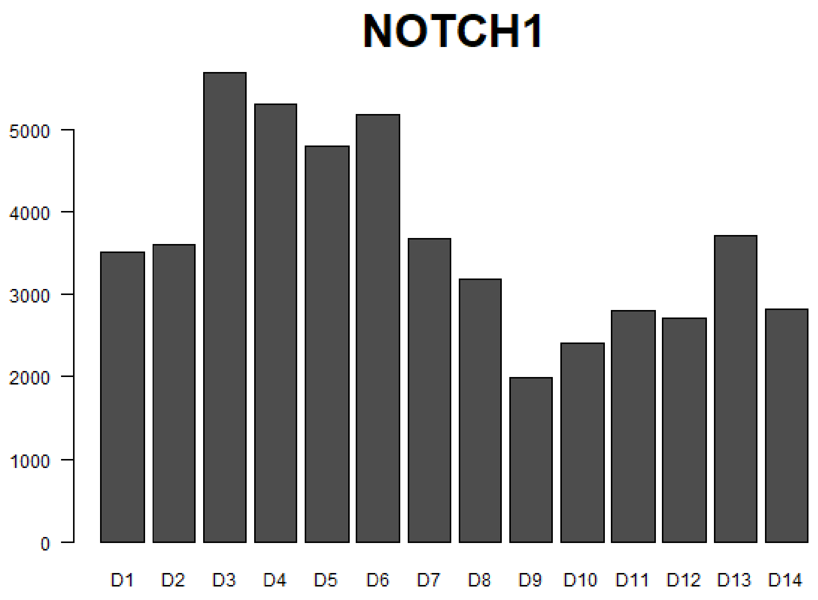


**Figure S19: The gene NOTCH1 is transiently over-expressed following release from cell cycle arrest and is associated with topic k2.**

The gene NOTCH1 was previously found to be over-expressed in cervical cancer with respect to normal cervical tissues (Sun et al. 2015; Zagouras et al. 1995).

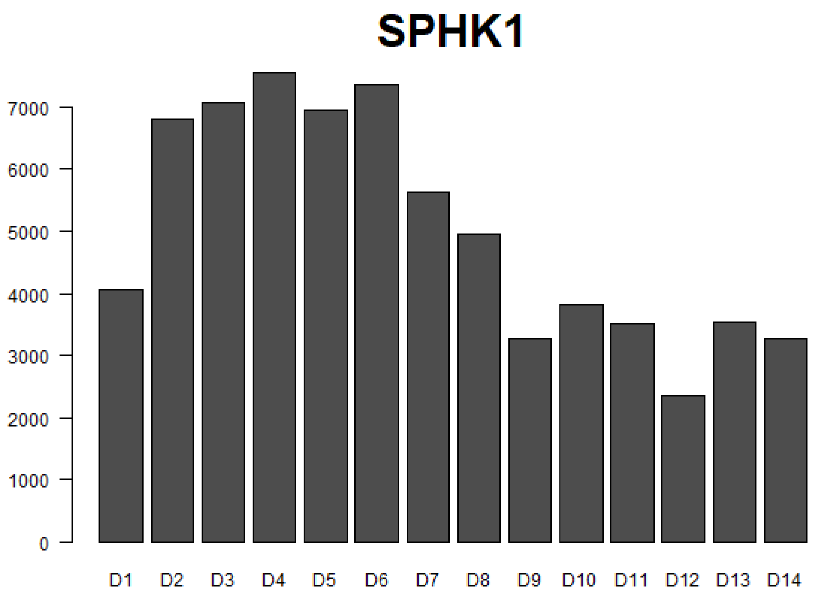


**Figure S20: The gene SPHK1 is transiently over-expressed following release from cell cycle arrest and is associated with topic k2.**

The gene SPHK1 is thought to promote inhibition of apoptosis and increased cell proliferation. This gene was previously found to be over-expressed in cervical cancer with respect to normal cervical tissue and to be associated with tumor stage, size, invasion, metastasis, and low survival (H.-S. Kim et al. 2015).

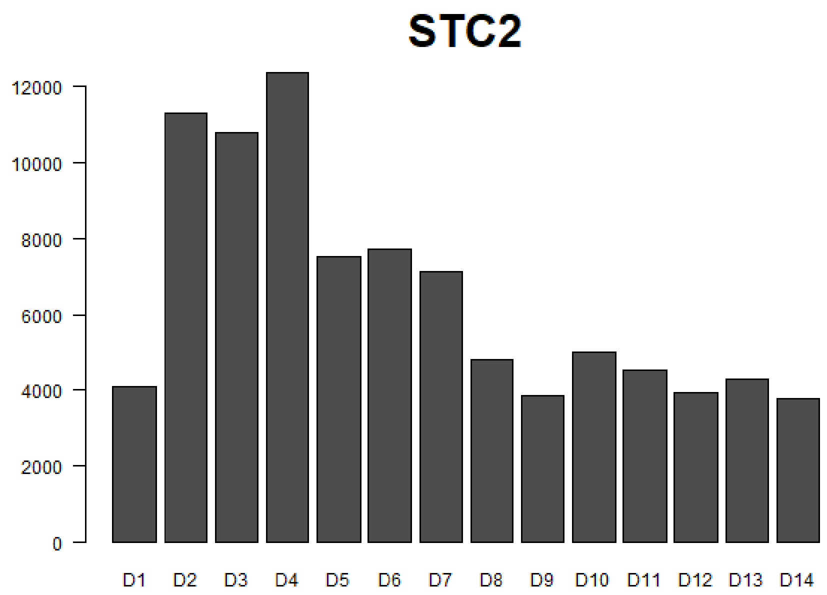


**Figure S21: The gene STC2 is transiently over-expressed following release from cell cycle arrest and is associated with topic k2.**

The gene STC2 was previously found to be significantly increased in cervical cancer tissues and cell lines compared to normal cervical tissues (Y. Wang et al. 2015) and to promote proliferation in cervical cancer cell lines, including HeLa cells.

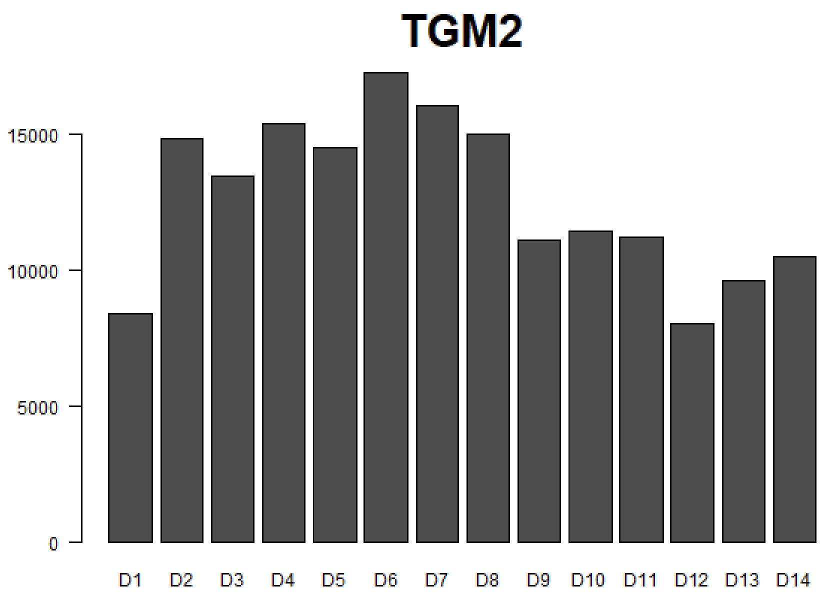


**Figure S22: The gene TGM2 is transiently over-expressed following release from cell cycle arrest and is associated with topic k2.**

The gene TGM2 was previously found to be over-expressed in cervical cancer relative to normal cervical samples (Caffarel et al. 2013).

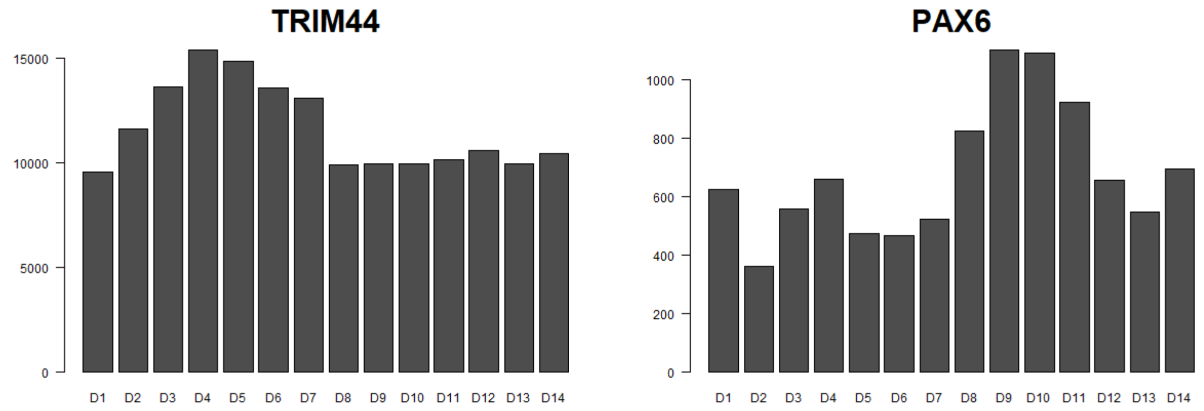


**Figure S23: The gene TRIM44 is transiently over-expressed following release from cell cycle arrest and is associated with topic k2.**

The gene TRIM44 was previously found to be significantly up-regulated in cervical cancer compared to normal cervical tissue, and also to be associated with increased tumor stage, grade, metastasis, and overall poor prognosis (Liu et al. 2019). Interestingly, we also observed that PAX6, whose expression is thought to be inhibited by TRIM44 (X. Zhang et al. 2015), becomes transiently over-expressed later, after TRIM44 over-expression has subsided (time points D8-D14).

.

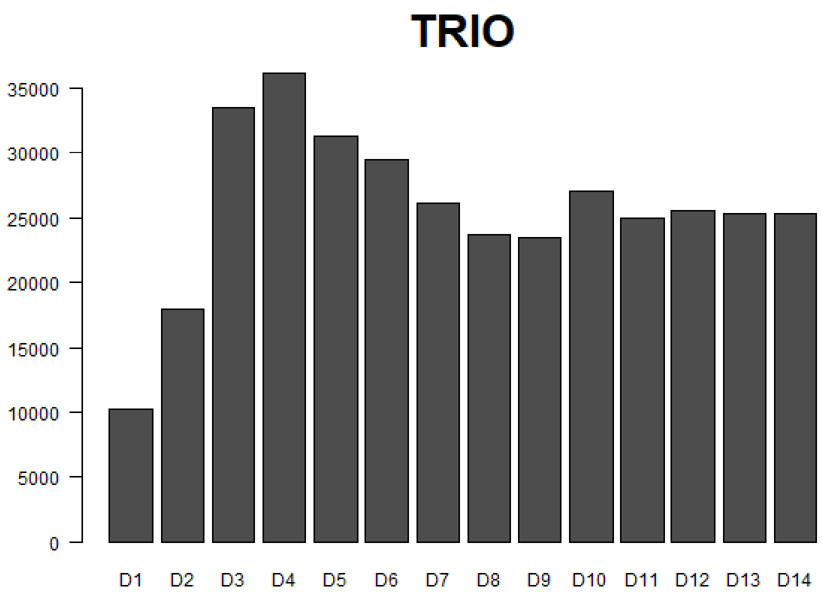


**Figure S24: The gene TRIO is transiently over-expressed following release from cell cycle arrest and is associated with topic k2.**

The gene TRIO was previously found to be over-expressed in cervical cancer with respect to adjacent normal tissues, and to be associated with cell migration (van Rijssel and van Buul 2012) and metastasis (Hou et al. 2018). Interestingly, we observed that during the first few hours after release from cell cycle arrest, the expression of this gene increases rapidly (time points D1-D4), 'overshoots', and then stabilizes during the second cell cycle (time points D8-D14).

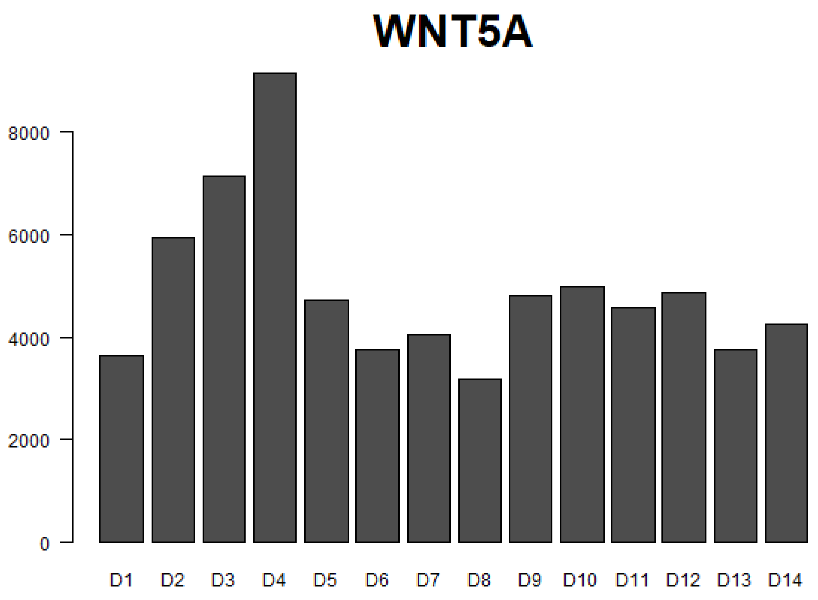


**Figure S25: The gene WNT5A is transiently over-expressed following release from cell cycle arrest and is associated with topic k2.**

The gene WNT5A was previously found to be overexpressed in cervical cancer compared to adjacent normal cervical tissues, and to be associated with tumor metastasis, recurrence, and shorter survival times (Lin et al. 2014).

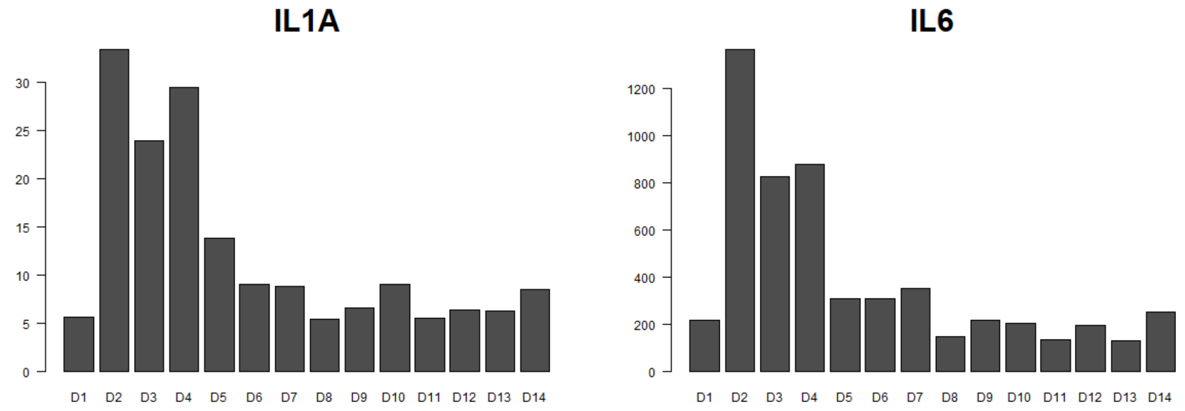


**Figure S26: The genes IL1A and IL6 are transiently over-expressed following release from cell cycle arrest and are associated with topic k2.**

The gene IL6 is an immediate-early response gene (Tullai et al. 2007). The genes IL1A and IL6 were both previously found to be over-expressed in cervical cancer compared to adjacent non-tumor tissue, and to be associated with cervical cancer progression and shorter survival times (Song et al. 2016).

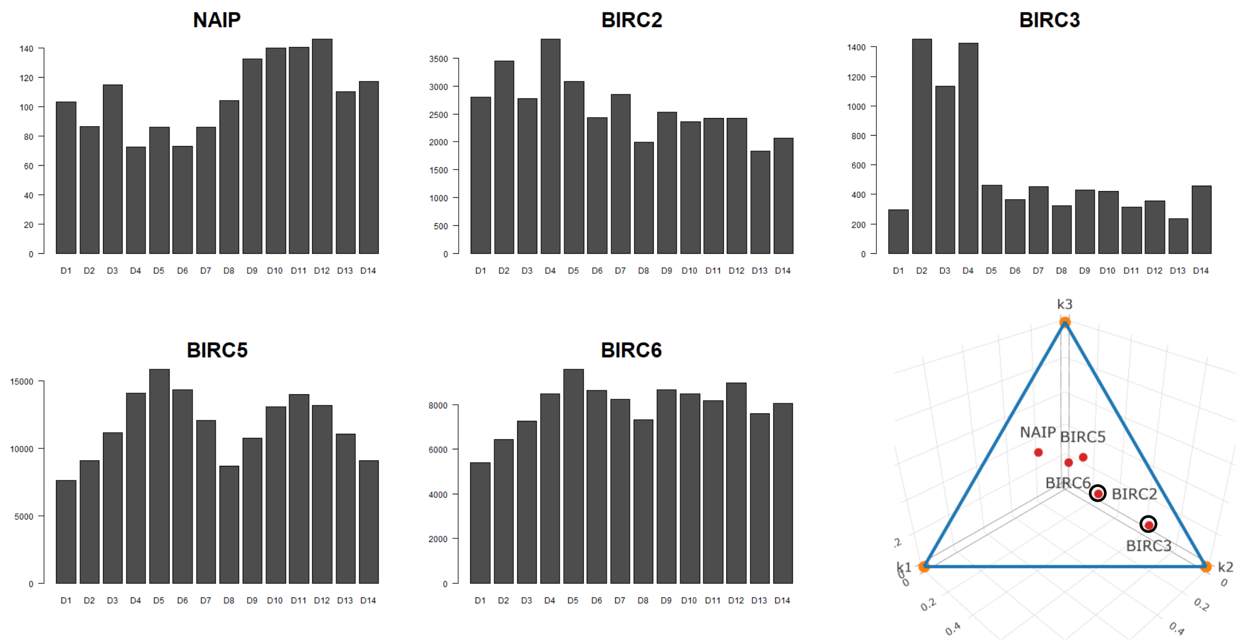


**Figure S27: The genes BIRC2 and BIRC3 are transiently over-expressed following release from cell cycle arrest and are associated with topic k2.**

The human BIR-containing protein family (BIRCs) is thought to contain two subgroups, each having structural and functional characteristics (Silke and Vaux 2001). The first subgroup includes the genes BIRC1 (NAIP), BIRC2, and BIRC3, which are presumed inhibitors of apoptosis, while the second group includes the genes BIRC5 (Survivin) and BIRC6 that are presumably regulators of the cell cycle and are required for mitotic chromosome segregation and cytokinesis. In the dataset that we studied we observed that the apoptosis inhibitors BIRC2 and BIRC3 are transiently over-expressed following release from cell cycle arrest, while the cell cycle regulator BIRC5, and to some extent also BIRC6, are periodically expressed. This can be seen also in the diagram of posterior probabilities (bottom right). Surprisingly, BIRC1 (NAIP) seems to be transiently repressed following release from cell cycle arrest. These differences in expression and function may be attributed to structural differences, for example, BIRC2 and BIRC3 both contain a caspase-recruitment domain (CARD) whereas BIRC1 (NAIP) has a nucleotide-binding loop instead (Silke and Vaux 2001).
